# *Bdellovibrio* and like organisms bacterial predators are not equally distributed in peri-alpine lakes

**DOI:** 10.1101/194688

**Authors:** Benoit Paix, Stéphan Jacquet

## Abstract

Microbes drive a variety of ecosystem processes and services but still many of them remain largely unexplored because of our lack of knowledge on both diversity and functionality of some potentially key microbiological compartments. This is typically the case with and within the group of bacterial predators collectively known as *Bdellovibrio* and like organisms (BALOs). Here we report for the first time the abundance, distribution and diversity of the three main families of these natural and obligatory predators of gram negative bacteria in three peri-alpine lakes (e.g. lakes Annecy, Bourget and Geneva) at different depths (surface *vs*. 45 or 50 m) and along a few months (from August 2015 to January 2016). We show that, using PCR-DGGE and cloning-sequencing approaches, the diversity appeared relatively low and very specific to fresh waters or even of the lakes themselves. While the *Peredibacteraceae* family was represented mainly by a single species (i.e. *Peredibacter starii*), it could constitute up to 7% of the total bacterial cell abundances. Comparati vel y, the abundances of the two other families (referred to as *Bdellovibrionaceae* and *Bacteriovaracaceae*) were significantly lower. More interestingly, the distribution in the water column was very different between the three groups suggesting various life strategies/niches for each of them: *Peredibactereacea* dominated near surface while the *Bdellovibrionaceae* and the *Bacteriovaracaceae* were more abundant at depth. All in all, our results suggest that these bacterial predators are likely to play a significant role in mortality, carbon fluxes and prokaryotic community structure in lakes.

## Introduction

Over the last few years, studies on western European large and deep peri-alpines lakes have shown that these ecosystems own a very diverse and dynamic auto- and heterotrophic prokaryotic community (Comte *et al.*, 2006 ; Personnic *et al.*, 2009 ; Debroas *et al*., 2009, Berdjeb *et al*., 2011 ; Berdjeb *et al*., 2013 ; Domaizon *et al.*, 2013 ; Ammini *et al.*, 2014). These studies and others have also revealed that a large variety of both biotic and abiotic factors are likely to regulate these communities. Among these factors, inorganic nutrients, viruses, nanoflagellates and other heterotrophic grazers (including ciliates and/or metazooplankto n) have been identified as key players in the dynamics of abundance or community composition structure patterns (Domaizon *et al*., 2003 ; Comte *et al*., 2006 ; Personnic *et al.*, 2009b ; Thomas *et al.*, 2011 ; Berdjeb *et al*., 2011; Berdjeb *et al*., 2013; Perga *et al*., 2013). Clearly, viral lysis and nanoflagellate or ciliate grazing have been observed as important biotic factors involved in bacterial mortality, affecting their abundance with a rate ranging from 10 to 60 % of bacterial loss per day, but also their structure and their diversity (Jacquet *et al*., 2005 ; Sime-Ngando *et al*., 2008 ; Thomas *et al.*, 2011 ; Meunier and Jacquet, 2015).

Other biotic interactions have been poorly investigated in these lakes. Among them the interactions between micro- and macroorganisms, the interactions between bacteria and other organisms or the role of eukaryotic pathogens (e.g. fungi) remain anecdotic (Humbert *et al*., 2009 ; Mangot *et al*., 2011 ; Perga *et al*., 2013). Comparatively to these or again the role of nanoflagellate or viruses mentioned above, another type of biotic interaction that has been largely neglected in aquatic ecosystems is bacterial predation by other bacteria and, to the best of our knowledge, the diversity, abundance, dynamics and functional role of this group of predators has never been investigated in peri-alpine lakes. Most of these predatory bacteria are grouped in *Bdellovibrio* and like organisms (BALOs). These bacteria are small, very motile, gram-negative and obligate predators that prey on other gram-negative bacteria. BALOs are widely distributed in different ecosystems like saltwater, freshwater, sewage, soils and sediments (Rotem *et al*., 2014 ; Perez *et al*., 2016). However, their abundance, their taxonomic diversity and the diversity of their habitats have been largely unexplored or underestimated because of the use of culturing approaches. The use of culture independent methods, using 16S DNA approaches, has revealed that the diversity of cultivated BALOs represents in fact only a small fraction of the actual diversity (Davidov *et al*., 2006).

*Bdellovibrio* and like organisms represent a polyphyletic group and can be found within two different classes: the *α*-proteobacteria with the genus *Micavibrio* and, merely, the *δ-*proteobacteria with three families: the *Bdellovibrionaceae*, the *Bacteriovaracaceae*, and the *Peredibacteraceae* (Rotem *et al*., 2014). The analysis of 16S rDNA sequences diversity in different environmental samples has revealed that *Bdellovibrionaceae* and *Bacteriovoracaceae* could represent the most diversified groups of BALOs in the environment. *Bdellovibrionaceae* are mostly represented by *Bdellovibrio bacteriovorus*, a periplasmic predator and *Bdellovibrio exovorus*, an epibiotic predator. They can be isolated from a variety of environments (Davidov & Jurkevitch, 2004). The *Bacteriovoracaceae*, that have been renamed *Halobacteriovoraceae,* are mainly represented by *Bacteriovorax litoralis* and *Bacteriovorax marinus*. They are generally associated to marine ecosystems (Baer *et al*., 2004 ; Piñeiro *et al*., 2008). *Peredibacteraceae* have been recently separated from the family formerly named *Bacteriovoracaceae* (Davidov & Jurkevitch, 2004). They are mainly represented by the species *Peredibacter starrii* and *Bacteriolyticum stolpii* (Rotem *et al*., 2014 ; Williams *et al.*, 2016) and are generally associated to only fresh waters. At last, the genus *Micavibrio* represents a minor group within BALOs and is represented by *M. admirantus* and *M. aeroginovorus* which are both epibiotic predators (Rotem *et al*., 2014).

BALOs have been reported to play an important role in bacterial ecology by shaping the bacterial community (Lambina *et al*., 1987 ; Yair *et al*., 2003). However, the understanding of the ecology of this bacterial community remains largely unknown in many aquatic environments, especially natural systems, like large and deep lakes, for which there is almost no data yet. Interestingly, the work of Roux *et al*. (2012) in Lake Bourget, followed by the study of Zhong *et al*. (2015) for Lakes Annecy and Bourget (France), showed that there is an important single stranded DNA virus community in these lakes, the Microviridae which are abundant and diverse. This community presented a boom and bust dynamic, but the correlation between the abundance of these viruses and the abundance of total heterotrophic bacteria remained difficult to establish (Zhong *et al.*, 2015). However, some viruses of Microviridae are known to be parasitic of BALOs such as *B. bacteriovorus* (Brentlinger *et al*., 2002). Thus, the presence of a relatively abundant and diverse community of ssDNA viruses in peri-alpine lakes may suggest that there might also be an abundant and diverse community of cellular hosts, including the BALOs. These bacteria, by their potential trophic interactions with other communities of bacteria, could play a significant role in the functioning of the microbial compartment. It is therefore critical to study and understand the abundance, distribution and diversity of these bacterial predators as well as their environmenta l impact on the trophic network of these ecosystems (Perez *et al*., 2016; Williams *et al*., 2016).

Hence, the main objective of this work was to reveal for the first time the existence of BALOs in peri-alpine Lakes (Annecy, Bourget, and Geneva) and to try answering the following questions: (i) Can BALOs be easily detected in peri-alpine Lakes? (ii) What is the structure and diversity of the BALOs? (iii) What is the quantitative importance of the main groups among this community of predatory bacteria? (iv) Can we highlight relationships between the community of the BALOs and heterotrophic bacteria? (v) What environmental factors appear to be important in the regulation of these interactions?

## Material and methods

### Study sites and sampling strategy

Sampling was conducted at the reference station of the 3 largest natural deep lakes in France and Western Europe, i.e. Lakes Annecy, Bourget and Geneva. These ecosystems are characterized by different trophic status: mesotrophic for Lake Geneva, oligo-mesotrophic for Lake Bourget, and oligotrophic for Lake Annecy (Jacquet *et al*., 2012, 2014). The samples were taken at different depths, characteristics of the epi-, meta - and hypolimnion: 2 m, 15 m, 50 m and 100 m deep for Lakes Bourget and Geneva, and 3 m, 15 m and 45 m for Lake Annecy. These samples were taken on average once per month for each Lake (with the exception of Lake Annecy) on the period between August 2015 and January 2016, representing in all 60 samples. For each depth and sampling, 1 L of water was filtered successively through two types of 47 mm diameter polycarbonate filters, the first one of 2 μm (to obtain firstly the bacterial community attached to particles), and the second one of 0,2 μm (to retrieve only the so-called free-living bacteria). A total of 120 filters was prepared and kept at -20°C. Both physical and chemical parameters as well as total bacterial counts were obtained as reported in previous studies (Berdjeb *et al*., 2011; Zhong *et al*., 2013; Ammini *et al*., 2014; Jacquet *et al*., 2014). Physical, nutrients, chlorophyll *a* and other environmental parameters including total bacterial abundance using flow cytometry were obtained as previously described (Personnic *et al*., 2009; Thomas *et al*., 2011; Berdjeb *et al*., 2011).

### DNA extraction and PCR primers

From these filters, DNA extractions were conducted using the GenElute™ Bacterial Genomic DNA Kit. Different cultures of BALOs, referred to as HD100 and 109J for *Bdellovibrio bacteriovorus* and A3.12 for *Peredibacter starrii* (gently provided by Dr. E. Jurkevitch), used as positive controls for PCR assays (see below) were centrifuged (10 min, 4°C, 13000 g) in order to collect the pellet and extract the DNA. DNA concentrations were quantified and controlled in terms of quality using Nanodrop 1000 Spectophotometer and QUBIT 3.0 Fluorometer (Thermofischer Scientific) with three replicates for each sample.

Different primers for either PCR or qPCR were selected for their specificity to the 16S rDNA of the 3 BALOs families, i.e. the *Bdellovibrionaceae*, *Bacteriovoracaceae*, and *Peredibacteraceae*. A total of 12 primers (Table 1) were tested using different PCR and qPCR protocols. As no qPCR primers for quantifying *Peredibacteraceae* were available in any previous studies, we designed and tested new primers using NCBI/Primer-BLAST online tool (http://www.ncbi.nlm.nih.gov/tools/primer-blast/),FastPCR software (http://primerdigital.com/fastpcr.html) and MEGA6 software (http://www.megasoftware.net/). In that goal, 120 existing sequences of 16 rDNA of *Peredibacteraceae* were aligned using clustalW method with MEGA6. A consensus sequence was obtained and used to find specific primers with NCBI/Primer-BLAST. With a high stringency (i.e. the primer must have had at least 3 total mismatches to unintended targets, including at least 2 mismatches within the last 6 bps at the 3’ end ; targets that had 7 or more mismatches to the primer were ignored ; the target had a maximum size of 350 bps). Each primer couple designed was then verified by qPCR amplification and cloning sequencing.

**Table I.**
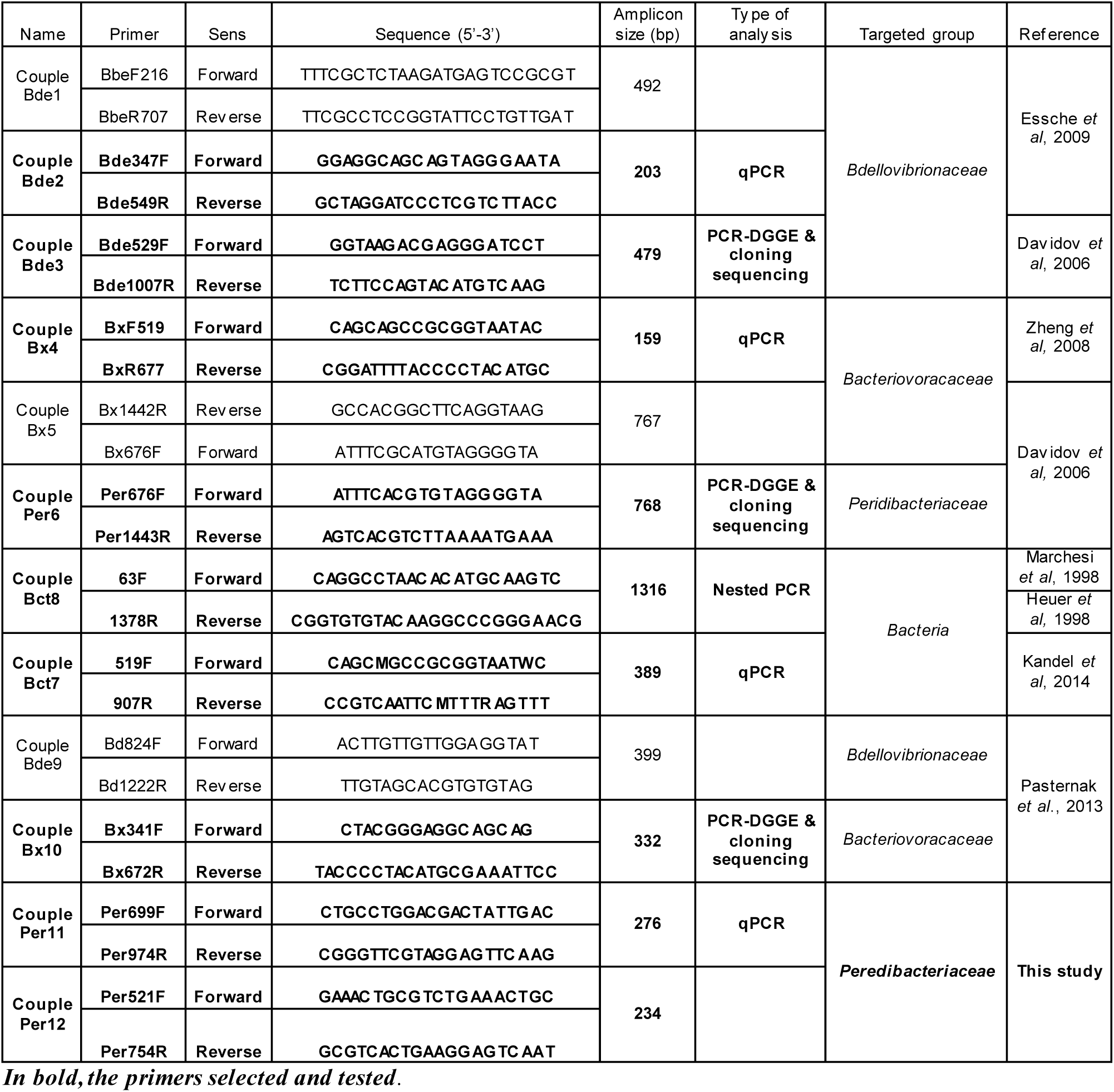
List of existing and selected primers used in this study

**Table II.** PCR conditions tested

| Primer names | Test | Tested PCR conditions |  |  |
| --- | --- | --- | --- | --- |
|  |  | Temp of hybridization | DNA volume. * | Number of cycles |
| Couple Bde1 | PCR | 49°C, 51°C, 55°C, 56°C, 58°C. | 1 and 2 µL | 36 cycles |
| Couple Bde2 | PCR | 49°C, 51°C, 55°C, 56°C, 58°C. | 1 µL | 36 cycles |
|  | qPCR | 60°C. | 1 µL | 45 cycles |
| Couple Bde3 | PCR | 49°C, 51°C, 55°C, 56°C, 58°C. | 1 and 2 µL | 36 cycles |
| Couple Bde9 | PCR | 48°C, 52°C. | 1 µL | 36 cycles |
| Couple Bx4 | PCR | 49°C, 54°C, 56°C, 58°C. | 1 µL | 36 cycles |
|  | qPCR | 58°C, 60°C. | 1 and 2 µL | 45 cycles |
| Couple Bx5 | PCR | 49°C, 51°C, 55°C, 56°C, 58°C. | 1 and 2 µL | 36 cycles |
| Couple Bx10 | PCR | 48°C, 52°C. | 1 µL | 36 cycles |
| Couple Per6 | PCR | 49°C, 51°C, 55°C, 56°C, 58°C. | 1 µL | 36 cycles |
| Couple Bct7 | PCR | 50°C. | 1 µL | 36 cycles |
|  | qPCR | 52°C. | 1 µL | 30, 40 & 45 cycles |
| Couple Bct8 | PCR | 48°C, 50 and 52°C. | 1 µL | 36 cycles |
| Couple Bct8<br>+ Couple Bde3 | Nested PCR 1 | Couple 8 : 50°C. | 1 µL | 20 cycles |
|  | Nested PCR 2 | Couple 3 : 51°C. | 1 µL | 20 cycles |
| Couple Bct8<br>+ Couple Bx10 | Nested PCR 1 | Couple 8 : 50°C. | 1 µL | 20 cycles |
|  | Nested PCR 2 | Couple 10 : 54°C. | 1 µL | 20 cycles |
| Couple Per11 | PCR | 53°C | 1 and 2 µL | 36 cycles |
|  | qPCR | 53°C | 1 and 2 µL | 45 cycles |
| Couple Per12 | PCR | 53°C | 1 and 2 µL | 36 cycles |

qPCR reactions were performed using the QuantiTect SYBR® Green PCR Kit and with the Rotor-Gene Q thermocycler. Standard curves were established in triplicates using serial dilutions of *E.coli* plamids containing 16S rDNA sequences of each three families. Linear standards curves were obtained within the range of 10^1^ to 10^6^ plasmid copies per reaction. The efficacy was 0.99 with a R^2^ value of 0.998 and a slope value of -3.32. The specificity of reactions was confirmed by both melting-curve analyses and agarose gel electrophoresis to identify unspecific PCR products. The plasmid copy numbers were calculated using the following formula (Whelan *et al*., 2003): Copy number = (DNA amount (ng) × 6.022 × 10^23^) / (length (bp) × 109 × 650).

### DGGE

The BALOs community was analyzed by denaturing gradient gel electrophoresis (DGGE) following the manufacturer’s protocol Instruction Manual (C.B.S.-Scientific company.inc, DGGE-2001). One mm thick polyacrylamide gel (6% [wt./vol] acrylamide in 1 × TAE buffer [40 mM Tris, 20 mM sodium acetate, 1 mM EDTA]; pH adjusted to 7.4) was prepared with a linear formamide/urea gradient ranging from 40% to 55% after several tests to find the best gradient. It was overlaid with a non-denaturing stacking gel. Each well was loaded with 15 ng PCR product and 5 μL loading buffer. Electrophoresis was conducted for 16 h at 120 V and 60°C. Subsequently, the gels were stained in darkness for 40 min in 1 × TAE buffer with 2 × SYBR gold solution as specified by the manufacturer. The DGGE profiles were only analyzed visually, because of the low number of bands obtained and the lack of apparent important diversity. Each band was cut under UV with Gel Doc™ XR+ system (Bio-Rad) and conserved with 30 μL of TAE buffer at -20°C. DNA extraction from DGGE band was performed by incubating tubes 20 min at -80°C, 20 min at -4°C and then by a centrifugating tubes (10 min, 13000 rpm, 4°C). The supernatant was conserved at 4°C for PCR.

### DNA purification, cloning and sequencing

The DNA of each DGGE band was eluted from the gel slice, after its excision, by adding 100 μl sterile 1 X TAE buffer and heating at 95°C for 15 min. Three microlitres of eluted DNA served as template in a 22 μl PCR mixture using the corresponding primer set. The PCRs were performed with the same conditions as the first PCR stage described above. The amplicons were first verified by electrophoresis in a 1.5% agarose gel, then purified using the I**l** ustra™ GFX™ PCR DNA and Gel Band Purification Kit (GE Healthcare), to finally be cloned into pCR®4-TOPO® vectors using the TOPO TA Cloning® Kit (Invitrogen). Randomly selected clones were sent for sequencing to GATC Biotech (Germany).

### Phylogenetic analysis

After cleaning and correcting sequences using BioEdit 7.0.5.3 (Hall, 1999), we obtained sequences have been deposited in GenBank under the following reference accession numbers XXX (this is in progress). All sequences were translated to amino acids, and aligned with culture representatives using MAFFT version 6 (Katoh *et al*., 2002). Multiple alignments were then curated using Gblocks (Castresana, 2000), employing a less stringent option that allowed for gaps inside the final blocks. We constructed the phylogenies using both the Bayesian inference and maximum likelihood methods. Baysian inference was conducted using MrBayes 3.2.1 (Ronquist *et al*., 2012) with two runs, four chains, 10^6^ generations, sampling every 100 generations, a burn-in value of 25%, and mixed models of amino acid substitution. The maximum likelihood phylogeny was constructed using PhyML 3.0 (Guindon *et al*., 2003), with 100 bootstrap replicates, and with the best models of acid-amino substitution and rate heterogeneity. The best models for each aligned-sequence-data-set were determined using MEGA 5 (Tumura *et al.*, 2011).

### Comparative analysis of taxonomic data

Some nMDS analyses were conducted to compare the consolidation of all of the sequences obtained and retrieved, in connection with the different types of ecosystems studied. The analysis is based on the construction of a matrix of dissimilarity between sequences (Bray-Curtis matrix), made from the Mothur software. Data issued from this matrix data were analyzed using R to generate a two-dimension nMDS from the vegan and VO packages. Statistical analyses were conducted with R realizing several PERMANOVA and repeated to check the significance (from the "p-value") of the grouping of sequences in different clusters.

### Data analysis of abundance in relation to environmental data

A principal component analysis (PCA) was made taking into account all available ecological monitoring environmental data to highlight the relationships or lack of relations between the abundances of various predatory and total bacteria (obtained by qPCR) with environmental factors. We built a matrix formed by 32 lines (3 lakes, the different dates and the two depths) and 14 columns (with the following environmental parameters: temperature, total phosphorus, orthophosphates, nitrates, ammonium, dissolved oxygen, chlorophyll *a*, pH, conductivity) and it was analyzed using XLStats™and R (Factoextra package), both of which gave similar results.

## Results

### Primer selection

Among the 12 primer sets tested by PCR or qPCR and checked using cloning-sequencing, we could choose one primer set for the phylogenetic analysis and another one for qPCR analysis, for each BALOs family (Tables 1 & 2). All selected primers were highly specific since all sequences obtained were characterized by more than 96% of identity with different cultured or uncultured bacteria of BALOs family found in databases (not shown). Among the two primers we designed to be able to quantify 16S rDNA sequence of the *Peredibactereceae* family by qPCR, Per699F (CTGCCTGGACGACTATTGAC) - Per974R (CGGGTTCGTAGGAGTTCAAG) was the best.

### Abundances and distribution of the BALOs

With a first analysis of absolute abundance of the different BALO families (in copies per mL, measured by qPCR) as a function of the different sampling months for the whole sampled water column without discrimination of depth, the data revealed that the most represented family of BALOs in the three lakes corresponded to the *Peredibacteraceae*, with an abundance up to 1.62 x 10^5^ copies per mL. In contrast, *Bdellovibrionaceae* and *Bacteriovoracaceae* were on average 10,000 times lower in abundance than the *Peredibacteraceae*, with maximum concentrations reaching barely 4 and 1.25 x 10^1^ copies per ml, respectively. Compared to total bacteria also quantified using qPCR, the *Peredibacteraceae* could represent up to 7.12% of total bacteria while *Bacteriovoracaceae* and *Bdellovibrionaceae* accounted for less than 0.05% of the bacterial community. Our results also revealed that the highest concentrations were always recorded in the free-living bacterial fraction suggesting that these BALOs were not attached bacteria. No clear seasonal variation was noticed here. When discriminating the three families at two distinct depths, i.e. the surface (2 m, 2.5 m or 3 m depending on the lake) vs. deeper waters (45 m or 50 m depending on the lake), (i) *Peredibacteraceae* were the most abundant (with 100 to 10000 times more copies per mL than *Bdellovibrionaceae* and *Bacteriovoracaceae*), (ii) *Peredibacteraceae* were generally more abundant at surface compared to depth while we observed the reverse for *Bdellovibrionaceae* and *Bacteriovoracaceae*. This contrast was particularly noticeable for Lake Bourget (Figure 1).

**Figure 1.**
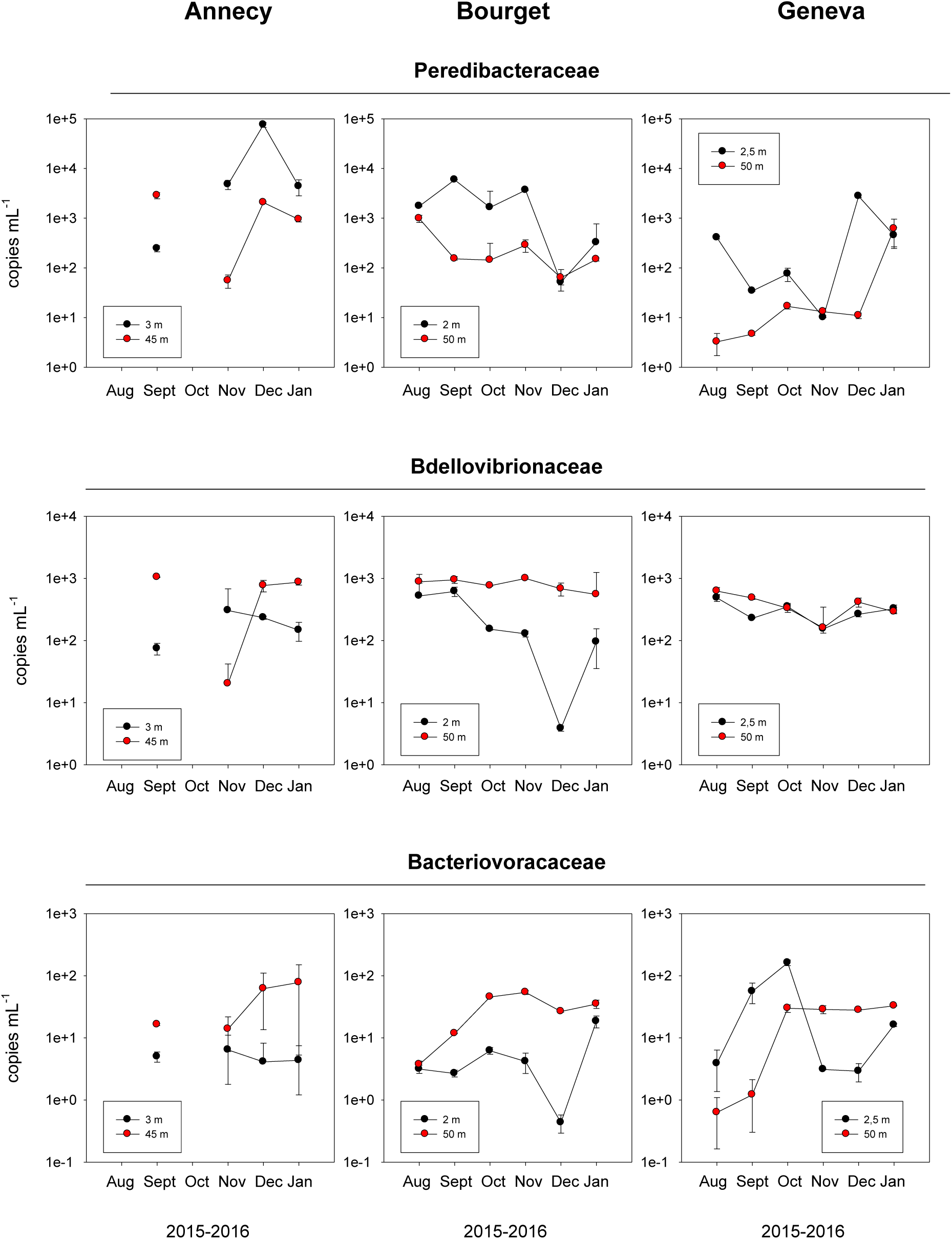
Dynamics of abundance for the different BALOs obtained at two contrasting depths in the three lakes

### Relationships between BALOs and total bacteria and other environmental data

Using abundance (from both qPCR and flow cytometry) and environmental data obtained from the *in situ* survey of peri-alpine lakes (e.g. http://www6.inra.fr/soere-ola/), ACP could be conducted to assess possible relationships between BALOs and their biotic and abiotic environment (Figure 2A & 2B). First, a positive and significant correlation (r=0.74, p<0.05) was observed between total heterotrophic bacteria and *Peredibacteraceae*, especially for surface waters of Lake Annecy. *Bdellovibrionaceae* displayed clear links with conductivity (r=0.44, p<0.05), oxygen (r=-0.48, p<0.05) and chlorophyll *a* (r=0.5, p<0.05) concentrations. Comparatively to the two other families, there was no significant relationship found for the *Bacteriovoracaceae*. Moreover, the two distinct water layers, i.e. the surface (<3 m depending on the lake) vs. deeper waters (>45 m depending on the lake), could be separated (Figure 2A). This analysis showed that *Bdellovibrionaceae* and *Bacteriovoracaceae* are more abundant in deep water and driven by ecological factors specific of deep waters. By contrast, *Peredibacteraceae* were more abundant in surface waters and driven by ecological factors more specific to surface waters (such as a higher concentration of total bacteria, dissolved O^2^ chlorophyll *a*, and a higher temperature). Also, samples could be separated into three distinct groups corresponding of each lake (Figure 2B). *Bacteriovoracaceae* seemed to be preferentially driven by ecological variables shared by Lake Geneva and Bourget (Mesotrophic and oligo - mesotrophic lakes), and *Bdellovibrionacace* by ecological variables shared by Lake Bourget and Annecy (Oligo-mesotrophic and oligotrophic lakes). By comparison, *Peredibacteraceae* was mainly associated to Lake Annecy characteristics.

**Figure 2.**
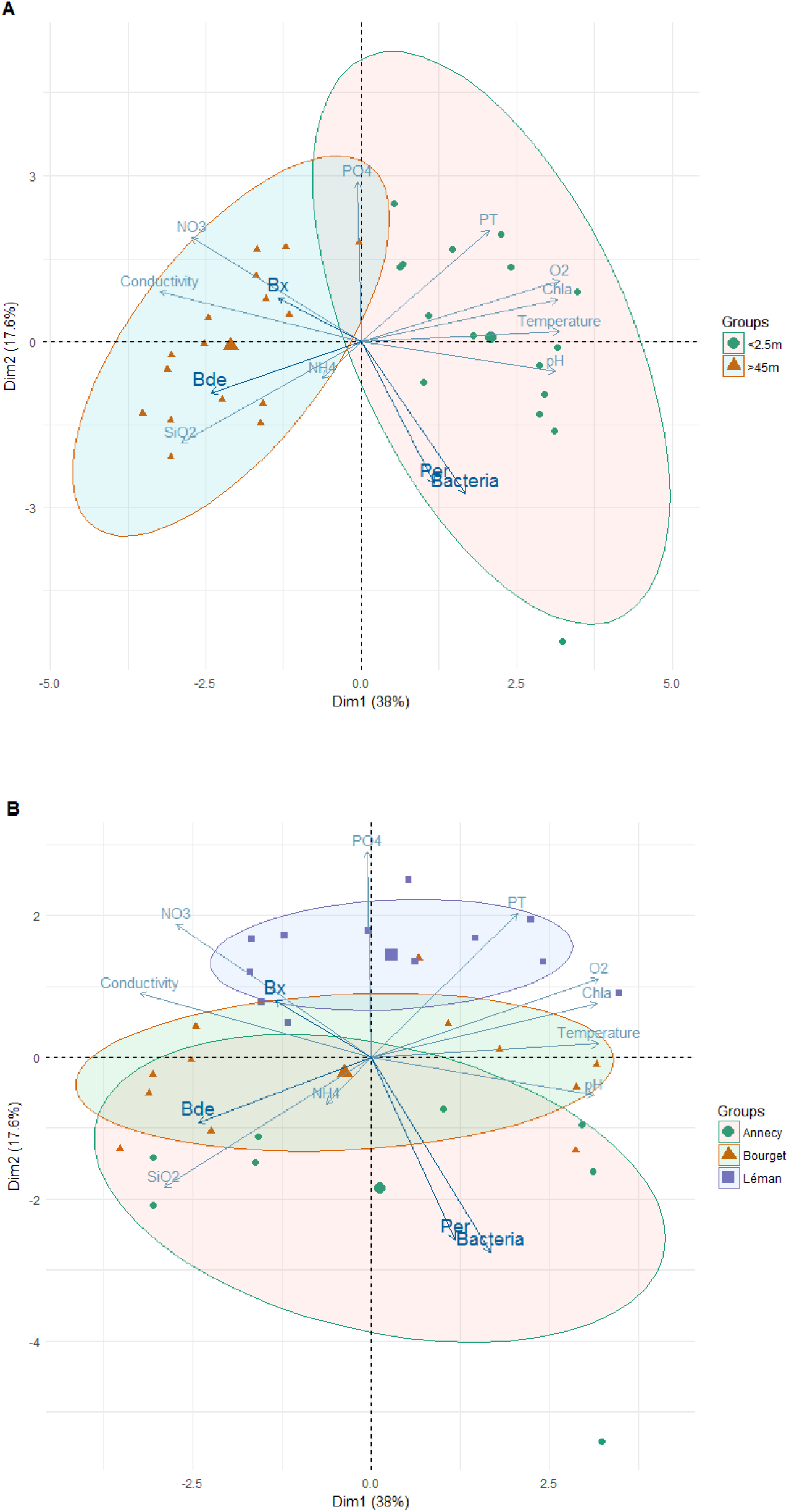
Principal Component Analysis (PCA) of the distribution of absolute abundance of each group (“Bx” for Bacteriovoracace ae, “Bde” for Bdellovibrionaceae, “Per” for Peredibacteaceae, and “Bacteria” for total Bacteria) quantified by qPCR, according to ecological variables measured during samplings. (a) repartition of each absolute abundance grouped between the two distinct depths (surface vs. deeper waters), (b) repartition of each absolute abundance grouped between each lake (Lake Annecy, Lake Bourget, Lake Geneva)

### Structure of the different BALOs

The DGGE analysis revealed only a limited number of bands whatever the BALOs family and no seasonal pattern was recorded. Only 1 to 5 major bands could be detected, suggesting a low genotypic diversity. A maximum of three bands was detected for *Bacteriovoracaceae*. One major band with minor (in terms of intensity) bands was observed for *Bdellovibrionaceae*. At last, two bands were generally observed for *Peredibacteraceae* (supplementary figure 1).

### Diversity of the different BALOs and ecological repartition

A cloning-sequencing approach was chosen to assess the genetic diversity within the group of BALOs. The phylogenetic trees and nMDS analysis were constructed from a total of 116 sequences coding the 16S rDNA specific sequence of each BALOs (37 for the *Peredibacteraceae*, 40 for the *Bdellovibrionaceae* and 39 for the *Bacteriovoracaceae*). Our results clearly show (in agreement with the DGGE results) that the diversity of each BALO family was relatively low. We also reveal the existence of few different clusters characterized by percentages of similarities of sequences greater than 99%, within each tree of the three families. The phylogenetic tree representing the diversity of *Bdellovibrionaceae* (supplementary figure 2), shows the presence of 5 clusters of sequences obtained (with more than 99% similarity of sequences within each cluster). Two OTUs were clearly defined on this tree from 97% similarity of sequences. The first OTU grouping two cluster of sequences (cluster 1 and 2; 25 sequences in total) appears to be mainly affiliated with *Bdellovibrio bacteriovorus*, a periplasmic predator. The second OTU grouping mainly clusters 3, 4 and 5; 11 sequences in total) seems affiliated with *Bdellovibrio exovorus*, an epibiotic predator. nMDS analysis (Figure 3B) showing the distribution of the different sequences of *Bdellovibrionaceae* coming from this tree as a function of a distance calculated by a dissimilarity matrix, confirms that our sequences are grouped significantly in two OTUs. OTU 1, which is mainly affiliated with *Bdellovibrio bacteriovorus* and OTU 2 mainly associated with *Bdellovibrio exovorus* (r = 0.48, p <0.001). It also appears that the diversity observed seems to be structured homogeneously without significant distinction according to the three large alpine lakes (p>0.05). Moreover, it can be highlighted that the sequences of *Deltaproteobacteria* which are not belonging to the *Bdellovibrionaceae* are clustered independently to the rest of the *Bdellovibrionaceae* sequences. The phylogenetic tree representing the diversity of *Bacteriovoracaceae* (supplementary figure 3) shows a relatively restricted diversity of sequences with clusters clustered at the base of the *Bacteriovoraceae* tree. The sequences obtained are grouped into different clusters that do not belong to any of the reference sequences found in NCBI, which mainly come from marine origin. This result suggests that the *Bacteriovoracaceae* sequences obtained in our study are still predominantly unaffiliated to a cultured *Bacteriovorax*, and thus are still unknown to date. The nMDS analysis (Figure 3C) based on the phylogenetic distance between each sequence confirms this result showing that sequences of marine origin and lacustrine origin constitute distinct clusters. Similarly to the phylogenetic analysis, nMDS analysis shows that the sequences obtained are grouped significantly into different distinct clusters (r = 0.63, p <0.001). These different clusters seem to correspond to the clusters observed on the phylogenetic tree. Moreover, no distinction between the sequences of the different clusters according to the lakes is observable nor significant (p> 0.05). Finally, the phylogenetic tree representing the diversity of *Peredibacteraceae* (supplementary figure 4) shows that the sequences obtained are taxonomically very close to 16S rDNA sequences of *Peredibacter starrii* (with a 99% similarity between them) which is represented by the strain *Peredibacter starrii* A3.12, isolated from fresh water (Pineiro *et al*., 2008). However, no sequence was directly associated to *Bacteriolyticum stolpii*, whose strain has also been isolated in fresh water (Pineiro *et al*., 2008). These results are supported by the nMDS analysis (Figure 3A) showing the significant taxonomic distribution of the 16S rDNA sequences preferentially affiliated with *Peredibacter starrii*. (r = 0.57, p <0.001). As for the two other families, no distinction is clearly visible, nor significant, regarding the taxonomic distribution of sequences obtained in relation to each lake (p <0.05).

**Figure 3.**
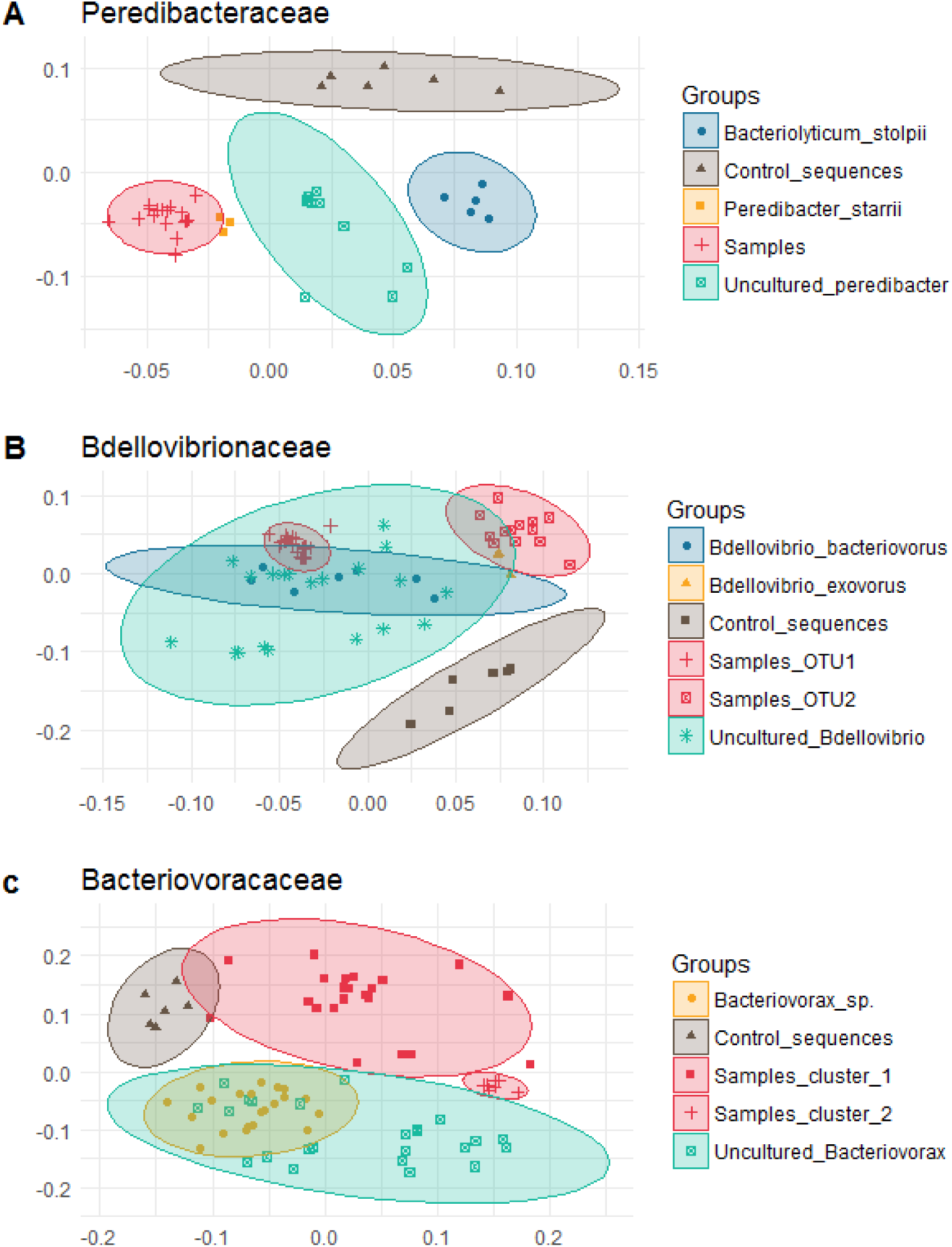
non-Metric MultiDimensional Scaling (nMDS) representing the distribution 16S rDNA sequences of each BALOs families obtained by cloning-sequencing method from samples, and 16S rDNA from NCBI databse. (a) Repartition of each 16S rDNA sequences of *Peredibacteraceae.* (b) Repartition of 16S rDNA sequences of *Bdellovibrionaceae*. (c) Repartition of each 16S rDNA sequences of *Bacteriovoracaceae*. For each nMDS, a same control group with 16s rDNA sequences of ?-proteobacteria which do not belong to any group of BALOs was used to verify the specificity of each 16s rDNA sequences from our samples.

## Discussion

The main result of this study is the revelation of the existence of specific bacterial predators (BALOs) in three large and deep French and Western Europe alpine lakes. Since the presence and importance of BALOs were examined in these models of freshwater ecosystems for the first time, different methods were tested and optimized in order to appreciate their diversity and abundance. We also had to design a new specific qPCR primer set for *Peredibacteraceae*, since no primer existed yet for this recently discovered freshwater BALO family. At last, we could quantify and assess the diversity of three representative groups of BALOs referred to as *Peredibacteraceae*, *Bdellovibrionaceae* and *Bacteriovoracaceae*, and compare these results to those obtained from a variety of other environments.

Even if our study clearly highlighted the existence and the relative importance of BALOs in the large peri-alpine lakes, it remains limited by the choice of methods used and the limited number of samples analyzed. We only worked with a few depths and for a few months, resulting to a preview of the distribution and dynamics of this community which is necessarily limited. This may partly explain why we were unable to clearly reveal key factors affecting the abundance or diversity of these predators. Regarding genetic analysis, we found only a fairly limited diversity. It is possible that the primers used were only able to amplify a part of the diversity within each family, and in a different way between the different groups. We are aware of the limitations associated with the conventional cloning-sequencing method based on DNA extraction of DGGE bands. Our results may have been limited and it is therefore not excluded that the diversity of the BALOs is in fact much higher. Therefore, further studies must use Next Generation Sequencing with different BALOs specific primers to better assess their diversity.

While *Peredibacteraceae* may be not a much diversified family, these bacteria were found in very high concentration compared to *Bdellovibrionaceae* and *Bacteriovoracacea*. *Peredibacteraceae*, whose strains described in the literature were isolated from freshwater (Pineiro *et al*., 2008), constituted the most abundant family for all the conditions studied (within the three lakes, depths, different fractions and different sampling periods). This first result suggests that the abundance of BALOs in the three tested peri-alpine lakes was likely linked to the very low salinity of these ecosystems. A low sodium chloride concentration but also a relatively diverse and high concentration of bacterial preys could clearly favor the development of BALOs adapted to the freshwater environment such as *Peredibacter starii* and would be linked to a low abundance of bacteria adapted to marine environments such as *Bacteriovorax* sp.

Lower concentrations of *Bdellovibrionaceae* and *Bacteriovoracaceae* (up to 125 copies per mL), compared to *Peredibacteraceae*, were more significant when compared to other studies. The study of the abundance of *Bdellovibrionaceae* and *Bacteriovracaceae* in aquaculture systems establishes measured concentrations between 10^3^ and 10^6^ cells per mL (Kandel et al., 2014). These results suggest, at first sight, that these two families had likely a more limited impact on the community of heterotrophic bacteria in our lakes. However, it is noteworthy that recent studies have shown that a low abundance of BALOs is not necessarily directly correlated with a lower functional impact on the dynamics of their prey (Richards et al., 2012, Williams et al., 2016, Welsh et al., 2016). Hence, despite the very low abundances of *Bdellovibrionaceae* and *Bacteriovoracaceae* found in the context of this study, their functional role may be not necessarily negligible.

## Conclusions

Our results reveal for the first time in large and peri-alpine lakes the presence of bacterial predators belonging to the three main families of BALOs. These results, discussed above, lead to the conclusion that these bacteria undoubtedly play a significant role in the functioning of these ecosystems, but that this role remains to be fully understood, especially for *Peredibacteraceae*, due to their numerical majority but also for *Bdellovibrionaceae* and *Bacteriovoracaceae*, for their diversity. A first perspective to our work is to be able to appreciate finely the interactions between host and predators through approaches like NGS (to also appreciate more finely diversity) with specific primers of each family of BALOs (networks of interactions) and also like experimental approaches from isolates. An approach using isolates from different strains representative of the BALO families, based on the work carried out by E. Jurkevich and his team, would make it possible to co-culture these bacteria in microcosms, with their prey in order to know the influence of different environmental parameters on the prey-predator relationship. Experiments in micro- or mesocosms could, for example, be imagined under different conditions, following the experimental approach proposed by Williams et al. (2016) enabling the enumeration of BALOs by qPCR and by a SIP type approach following the addition of radiolabelled preys in different conditions. The enumeration and analysis of the diversity of the various BALOs in cosms would allow to directly correlate the different environmental parameters characteristic of the lacustrine environment (such as prey quantity, prey diversity, types of nutrients, etc.). …) with the distribution of the various BALOs. To these different perspectives proposed for the study of the functional role of BALOs within the water column, the importance of the abundance, structure and diversity of BALOs within other matrices, such as biofilms and sediments within the peri-alpine lake environment are another interesting issue. This would also involve re-examining the primers and perhaps designing new ones. Other questions remain unresolved, particularly regarding the diversity, the quantitative and functional importance of a family of BALOs not studied in this works: the *Micavibrio* group. Even if the identification of new species or BALO families remain largely unknown in peri-alpine lakes, data on marine, coastal or ocean environments are also very incomplete, and a study to compare BALOs in these different types of ecosystems would also be really interesting.

## Acknowledgments

We thank Dr. Edouard Jurkevitch who kindly provided us with the different BALOs strains and made a critical reading of the manuscript. We also thank Cécile Chardon for her help with molecular biology.

## References

Ammini P, Zhong X, Pradeep Ram AS, Jacquet S. (2014). Dynamics of auto- and heterotrophic picoplankton and associated viruses in Lake Geneva. Hydrol Earth Syst Sci 18: 1073–1087.

Baer ML, Ravel J, Piñeiro SA, Guether-Borg D, Williams HN. (2004). Reclassification of salt-water Bdellovibrio sp. as Bacteriovorax marinus sp. nov. and Bacteriovorax litoralis sp. nov. Int J Syst Evol Microbiol 54: 1011–1016.

Berdjeb L, Ghiglione J-F, Jacquet S. (2011a). Bottom-Up versus Top-Down Control of Hypo- and Epilimnion Free-Living Bacterial Community Structures in Two Neighboring Freshwater Lakes. Appl Environ Microbiol 77: 3591–3599.

Berdjeb L, Pollet T, Chardon C, Jacquet S. (2013). Spatio-temporal changes in the structure of archaeal communities in two deep freshwater lakes. FEMS Microbiol Ecol 86: 215–230.

Berdjeb L, Pollet T, Domaizon I, Jacquet S. (2011b). Effect of grazers and viruses on bacterial community structure and production in two contrasting trophic lakes. BMC Microbiol 11: 88.

Brentlinger KL, Hafenstein S, Novak CR, Fane BA, Borgon R, McKenna R, et al. (2002). Microviridae, a family divided: isolation, characterization, and genome sequence of phiMH2K a bacteriophage of the obligate intracellular parasitic bacterium Bdellovibrio bacteriovorus. J Bacteriol 184: 1089–1094.

Comte J, Jacquet S, Viboud S, Fontvieille D, Millery A, Paolini G, et al. (2006). Microbial community structure and dynamics in the largest natural French lake (Lake Bourget). Microb Ecol 52: 72–89.

Davidov Y, Friedjung A, Jurkevitch E. (2006). Structure analysis of a soil community of predatory bacteria using culture-dependent and culture-independent methods reveals a hitherto undetected diversity of Bdellovibrio-and-like organisms. Environ Microbiol 8: 1667–1673.

Davidov Y, Jurkevitch E. (2004). Diversity and evolution of Bdellovibrio-and-like organisms (BALOs), reclassification of Bacteriovorax starrii as Peredibacter starrii gen. nov., comb. nov., and description of the Bacteriovorax–Peredibacter clade as Bacteriovoracaceae fam. nov. Int J Syst Evol Microbiol 54: 1439–1452.

Debroas D, Humbert J-F, Enault F, Bronner G, Faubladier M, Cornillot E. (2009). Metagenomic approach studying the taxonomic and functional diversity of the bacterial community in a mesotrophic lake (Lac du Bourget – France). Environ Microbiol 11: 2412– 2424.

Domaizon I, Savichtcheva O, Debroas D, Arnaud F, Villar C, Pignol C, et al. (2013). DNA from lake sediments reveals the long-term dynamics and diversity of Synechococcus assemblages. Biogeosciences 10: 3817–3838.

Domaizon I, Viboud S, Fontvieille D. (2003). Taxon-specific and seasonal variations in flagellates grazing on heterotrophic bacteria in the oligotrophic Lake Annecy – importance of mixotrophy. FEMS Microbiol Ecol 46: 317–329.

Guindon S, Dufayard J-F, Lefort V, Anisimova M, Hordijk W, Gascuel O. (2010). New algorithms and methods to estimate maximum-likelihood phylogenies: assessing the performance of PhyML 3.0. Syst Biol 59: 307–321.

Humbert J-F, Dorigo U, Cecchi P, Le Berre B, Debroas D, Bouvy M. (2009). Comparison of the structure and composition of bacterial communities from temperate and tropical freshwater ecosystems. Environ Microbiol 11: 2339–2350.

Jacquet S, Domaizon I, Anneville O. (2012). Evolution de paramètres clés indicateurs de la qualité des eaux et du fonctionnement écologique des grands lacs péri-alpins (Léman, Annecy, Bourget): Etude comparative de trajectoires de restauration post-eutrophisation. Arch Sci 65: 225–242.

Jacquet S, Domaizon I, Anneville O. (2014). The need for ecological monitoring of freshwaters in a changing world: a case study of Lakes Annecy, Bourget, and Geneva. Environ Monit Assess 186: 3455–3476.

Jacquet S, Domaizon I, Personnic S, Ram ASP, Hedal M, Duhamel S, et al. (2005). Estimate s of protozoan- and viral-mediated mortality of bacterioplankton in Lake Bourget (France). Freshwater Biol 50: 627–645.

Kandel PP, Pasternak Z, van Rijn J, Nahum O, Jurkevitch E. (2014). Abundance, diversity and seasonal dynamics of predatory bacteria in aquaculture zero discharge systems. FEMS Microbiol Ecol 89: 149–161.

Lambina VA, Ledova LA, Churkina LG. (1987). Importance of Bdellovibrio in regulating microbial cenoses and self-purification processes in domestic sewage. Mikrobiologiia 56: 860– 864.

Mangot J-F, Debroas D, Domaizon I. (2011). Perkinsozoa, a well-known marine protozoan flagellate parasite group, newly identified in lacustrine systems: a review. Hydrobiologia 659: 37–48.

Meunier A, Jacquet S. (2015). Do phages impact microbial dynamics, prokaryotic community structure and nutrient dynamics in Lake Bourget? Biol Open 4: 1528–1537.

Pérez J, Moraleda-Muñoz A, Marcos-Torres FJ, Muñoz-Dorado J. (2016). Bacterial predation: 75 years and counting ! Environ Microbiol 18: 766–779.

Perga M-E, Domaizon I, Guillard J, Hamelet V, Anneville O. (2013). Are cyanobacterial blooms trophic dead ends ? Oecol 172: 551–562.

Personnic S, Domaizon I, Dorigo U, Berdjeb L, Jacquet S. (2009a). Seasonal and spatial variability of virio-, bacterio-, and picophytoplanktonic abundances in three peri-alpine lakes. Hydrobiologia 627: 99–116.

Personnic S, Domaizon I, Sime-Ngando T, Jacquet S. (2009b). Seasonal variations of microbial abundances and virus-versus flagellate-induced mortality of picoplankton in three peri-alpine lakes. J Plankton Res 31: 1161–1177.

Piñeiro SA, Williams HN, Stine OC. (2008). Phylogenetic relationships amongst the saltwater members of the genus Bacteriovorax using rpoB sequences and reclassification of Bacteriovorax stolpii as Bacteriolyticum stolpii gen. nov., comb. nov. Int J Syst Evol Microbiol 58: 1203–1209.

Richards GP, Fay JP, Dickens KA, Parent MA, Soroka DS, Boyd EF. (2012). Predatory Bacteria as Natural Modulators of Vibrio parahaemolyticus and Vibrio vulnificus in Seawater and Oysters. Appl Environ Microbiol 78: 7455–7466.

Rotem O, Pasternak Z, Jurkevitch E. (2014). Bdellovibrio and Like Organisms In: Rosenberg E, DeLong EF, Lory S, Stackebrandt E, Thompson F (eds). The Prokaryotes. Springer Berlin Heidelberg, pp 3–17.

Sime-Ngando T, Colombet J, Personnic S, Domaizon I, Dorigo U, Perney P, et al. (2008). Short-term variations in abundances and potential activities of viruses, bacteria and nanoprotists in Lake Bourget. Ecol Res 23: 851–861.

Thomas R, Berdjeb L, Sime-Ngando T, Jacquet S. (2011). Viral abundance, production, decay rates and life strategies (lysogeny versus lysis) in Lake Bourget (France). Environ Microbiol 13: 616–630.

Welsh RM, Zaneveld JR, Rosales SM, Payet JP, Burkepile DE, Thurber RV. (2016). Bacterial predation in a marine host-associated microbiome. ISME J 10: 1540–1544.

Williams HN, Lymperopoulou DS, Athar R, Chauhan A, Dickerson TL, Chen H, et al. (2016). Halobacteriovorax, an underestimated predator on bacteria: potential impact relative to viruses on bacterial mortality. ISME J 10: 491–499.

Yair S, Yaacov D, Susan K, Jurkevitch E. (2003). Small eats big: ecology and diversity of Bdellovibrio and like organisms, and their dynamics in predator-prey interactions. Agronomie 23: 433–439.

Zhong X, Berdjeb L, Jacquet S. (2013). Temporal dynamics and structure of picocyanobacteria and cyanomyoviruses in two large and deep peri-alpine lakes. FEMS Microbiol Ecol 86: 312– 326.

Zhong X, Guidoni B, Jacas L, Jacquet S. (2015). Structure and diversity of ssDNA Microviridae viruses in two peri-alpine lakes (Annecy and Bourget, France). Res in Microbiol 166: 644–654.

